# Parenteral glucose supply and pharmacological glycolysis inhibition determine the clinical fate of infected preterm newborns

**DOI:** 10.1101/2021.09.03.458815

**Authors:** Tik Muk, Anders Brunse, Nicole L. Henriksen, Karoline Aasmul-Olsen, Duc Ninh Nguyen

**Affiliations:** Section for Comparative Pediatrics and Nutrition, Department of Veterinary and Animal Sciences, University of Copenhagen, Denmark

**Keywords:** glycolysis, neonatal infection, neonatal sepsis, parenteral nutrition, preterm newborns

## Abstract

Preterm infants are susceptible to bloodstream infection that can lead to sepsis. High parenteral glucose supplement is commonly used to support their growth and energy expenditure, but may exceed endogenous regulation during infection, causing dysregulated immune response and clinical deterioration. Using a preterm piglet model of neonatal sepsis induced by *Staphylococcus epidermidis* infection, we demonstrate the delicate interplay between immunity and energy metabolism to regulate the host infection response. Circulating glucose levels, glycolysis and inflammatory response to infection are closely connected across the states of tolerance, resistance and immunoparalysis. Further, high parenteral glucose provision during infection induces hyperglycemia, elevated glycolysis and inflammation, leading to lactate acidosis and sepsis, whereas glucose restricted individuals are clinically unaffected with increased gluconeogenesis to maintain moderate hypoglycemia. Finally, pharmacological glycolysis inhibition during normoglycemia enhances bacterial clearance and dampens inflammation but fails to prevent sepsis. Our results uncover how blood glucose controls immune cell metabolism and function, in turn determining the clinical fate of infected preterm neonates. This also questions the current practice of parenteral glucose supply for infected preterm infants.

## Introduction

Millions of infants are born preterm (< 37 weeks of gestation) every year with up to 40% of them experiencing serious neonatal infection, leading to sepsis (1). Coagulase-negative Staphylococci (CONS) are responsible for up to 75% of nosocomial late-onset sepsis (LOS, > 3 days after birth) episodes with *S. epidermidis* being the predominant pathogenic species (2-4). Currently, there are no other effective therapies for neonatal infection than antibiotics, which are empirically used for almost all preterm infants despite only a small fraction of them indeed being infected (5, 6). Excessive antibiotic use predisposes to immunosuppression, secondary infection and antimicrobial resistance (7). Therefore, development of new infection therapies is of utmost importance.

Immunometabolism, the interplay between immune cell energy metabolism and function, has emerged as a key mechanism involved in many adult diseases, but its role in neonatal infection is unclear. Initial theoretical (8, 9) and *in vitro* reports (10, 11) suggest that the low energy reservoir in newborns programs the immune system to a disease tolerance strategy to avoid the Warburg effect in immune cells switching from oxidative phosphorylation (OXPHOS) to glycolysis, which quickly produces vast ATP amounts fueling inflammatory responses. This may explain how newborns, especially preterm newborns, can tolerate 10-100 times higher systemic bacterial loads (8) and have diminished blood cytokine responses to *in vitro* infection challenge (12, 13), relative to adults. However, it is still unclear how this disease tolerance of preterm infants is connected to their high susceptibility to neonatal sepsis, a pathological state characterized by an early hyper-inflammatory phase followed by immunoparalysis or death (14).

During the first few weeks of life, a majority of preterm infants receive parenteral nutrition (PN) to maintain sufficient nutrition, and international guidelines recommend high parenteral glucose supply (up to 17 g/kg/day) to avoid hypoglycemia (blood glucose <2.6 mM) and related brain injury (15-18). However, prolonged high parenteral glucose intake may lead to hyperglycemia (blood glucose >6.9 mM)(19), which is detected in up to 80% preterm infants (20). Notably, there are no specific guidelines for using parenteral glucose during neonatal infection, although PN-related hyperglycemia is associated with longer hospitalization in septic infants (16). We postulate that high parenteral glucose provision to infected newborns may accelerate blood immune cell glycolysis, driving excessive inflammation and leading to sepsis. Detailed understanding of this mechanism may shed light on novel therapies, e.g. reduced parenteral glucose supply or glycolysis inhibition.

Numerous animal infection and sepsis models have been established, e.g. cecal ligation and puncture (21), oral (22, 23) or systemic bacterial challenge (24). However, no rodent models can address the contributing effects of PN. The preterm pig is a unique model because it allows PN administration via umbilical catheter, similar to preterm infants (25). Further, preterm pigs mimic the immaturities of multiple organs and infection susceptibility in preterm infants (24, 26–28). Systemic *S. epidermidis* administration to newborn preterm pigs can induce clinical and cellular responses (fever, inflammation, immune cell depletion) progressing to septic shock (acidemia, and hypotension) 12-24h post-infection (24). Here, we further utilized this sepsis model and showed that the immunometabolic response to infection in preterm newborns was tightly regulated by circulating glycolysis-OXPHOS axis and glucose levels. We found that high parenteral glucose supply predisposed to hyperglycemia, excessive inflammation, reduced bacterial clearance and extreme sensitivity to sepsis following neonatal infection, while restricted parenteral glucose provision protected against sepsis. We also showed that a lesser reduction in glucose supply, with or without administration of a glycolysis inhibitor dichloroacetate (DCA), prevented hypoglycemia, enhanced bacterial clearance, alleviated systemic inflammation and lactic acidosis but did not protect against sepsis. Parenteral glucose restriction may be an effective and lifesaving therapy for infected preterm infants.

## Results

### *In vitro* and *in vivo S. epidermidis* thresholds determine the host immunometabolic responses

Preterm infants can presumably withstand higher circulating bacterial levels than adults and term infants prior to mounting resistant responses and later immunoparalysis (8). Here we first tested the threshold switching among those phases by measuring *in vitro* immunometabolic responses of cord blood from preterm pigs to increasing doses of *S. epidermidis* (Fig. 1A-F). At low bacterial doses (5×10^1^-5×10^4^ CFU/ml), inflammatory (TNFα) and anti-inflammatory (IL10) cytokine responses at both gene and protein levels were trivial, indicating an immune tolerant state (green, Fig.1 A-C, and Fig. S1A-C). At a dose of 5×10^5^ CFU/ml, a switch to resistant response occurred with an increase in TNFα and IL10 at both protein and gene levels (orange), relative to control and lower bacterial doses. Of note, the ratio of *TNFA*/*IL10* (Th1/Th2 cytokines) peaked at the dose of 5×10^5^ CFU/ml, but decreased again at higher doses, indicating another switch from resistant response to immunoparalysis (red). The same trends applied to other parameters, including elevated inflammatory targets (*IL*6, *TLR2*) and Th1 responses (*IFNG* and *IFNG*/*IL4*), and decreased regulatory T cell percentage at the bacterial dose of 5×10^5^ CFU/ml but not lower or higher doses (Fig. 1F and Fig. S1D-H). In parallel, cellular glucose uptake, measured by the differences in supernatant glucose levels with vs. without bacterial challenge, was gradually elevated with increasing bacterial doses, then reached a plateau level at the bacterial dose of 5×10^5^ CFU/ml (Fig. S1I). Further, genes related to OXPHOS (*COX1*) and glycolysis-mTOR pathway (*HIF1A*) were lowest and highest, respectively, also at the bacterial dose causing resistant responses (orange, Fig. 1D-E). These data revealed a clear dose-dependent switch of immunometabolic response to *S. epidermidis* from tolerance (low doses) to resistance (higher doses) and later immunoparalysis (very high doses).

**Figure 1.**
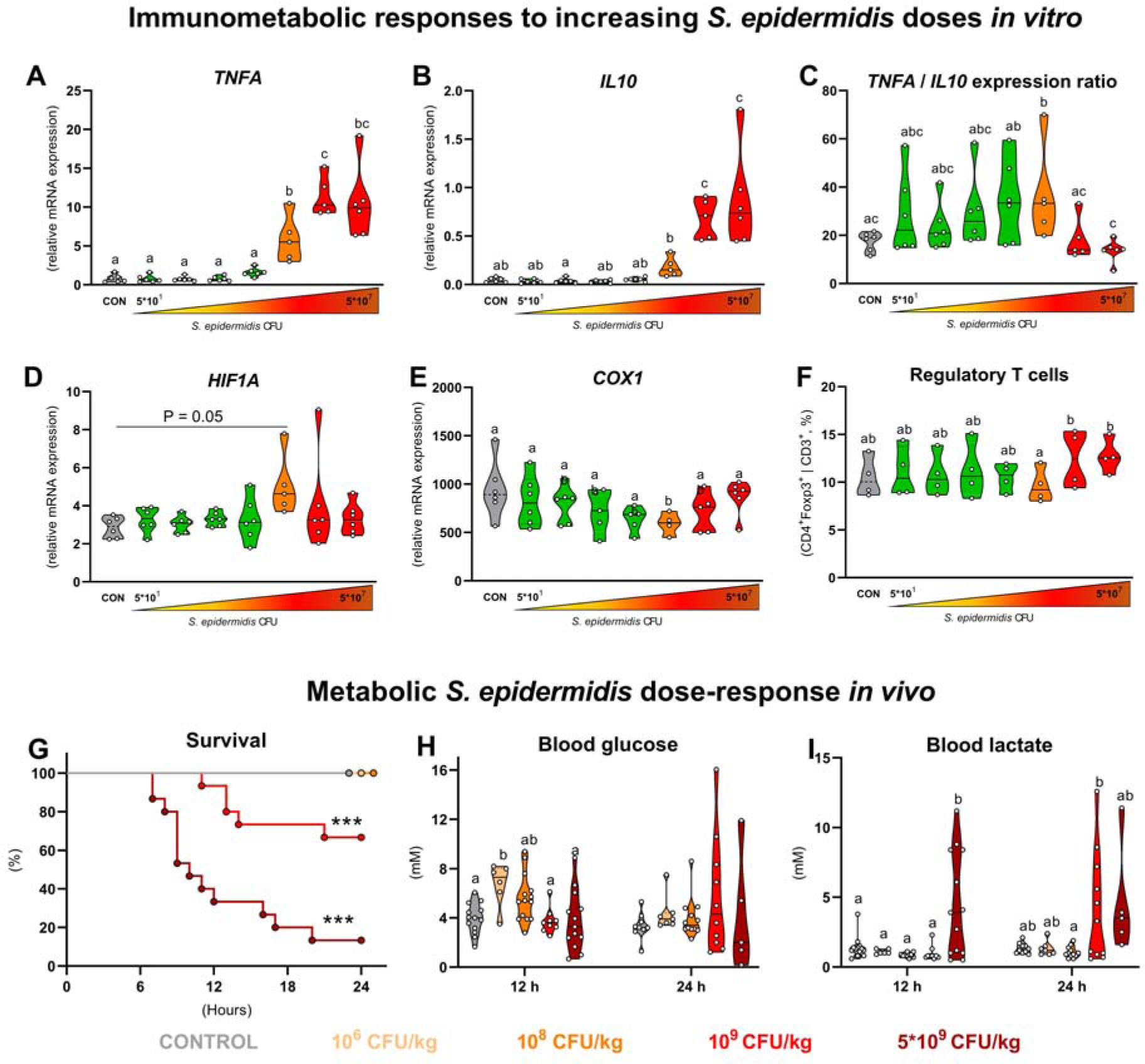
*In vitro* and *in vivo* immunometabolic response to *S.epidermidis*. (**A-E**) mRNA levels of *TNFA*, *IL10*, *TNFA*/*IL10* ratio, *COX1* and *HIF1A* of cord blood from preterm piglets in responses to an increasing bacterial dose (5×10^1^ -5×10^7^ CFU/mL, stimulated for 2 h at 37°C and 5%CO_2_, n = 5-6). (**F**) Frequency of Foxp3^+^ cells within the CD4^+^ lymphocyte population in *S. epidermidis*-stimulated cord blood (2 h at 37°C and 5%CO_2_, n = 4). (**G)** Survival rate, (**H**) blood glucose and (**I**) lactate levels of preterm newborn piglets 24 h post-infection with *S. epidermidis* (10^6^-5 x 10^9^ CFU/kg) via the intra-arterial catheter. Data in A-F, H-I are presented as violin dot plots with median (solid line) and interquartile range (dotted lines) and were analyzed using linear mixed-effect model followed by Tukey Post-hoc comparisons. *In vivo* data are presented as cumulative hazard curve or violin plots and were analyzed by Mantel-Cox test or linear model followed by Tukey Post-hoc comparisons. Values at a time point not sharing the same letters are significantly different (P < 0.05). *** P < 0.001, compared with the uninfected control.

We then tested clinical and metabolic responses to increasing *S. epidermidis* doses *in vivo*, using newborn preterm pigs (90% gestation) nourished by PN with a standard glucose level. The animals were clinically and metabolically unaffected by the two lowest doses (10^6^-10^8^ CFU/kg, disease tolerance). At a dose of 10^9^ CFU/kg, survival was 75% with dysregulated glucose and lactate at 24 h follow-up (disease resistance), whereas the highest dose of 5×10^9^ CFU/kg decreased 24 h survival to less than 20% and induced glucose and lactate dysregulation already at 12 h (Fig. 1G-I). Thus, clinical responses were clearly intertwined with perturbed glucose homeostasis and followed a severity spectrum dictated by *S. epidermidis* dose. Both *in vitro* and *in vivo* studies showed that immune cells had a propensity to undergo a metabolic shift towards aerobic glycolysis when activated, whereby glucose availability determined the potency of the cellular response with potential clinical implications.

### PN glucose determines sepsis susceptibility during *S. epidermidis* infection

We next investigated clinical, metabolic and immune responses to bacteremia on the background of extreme differences in glucose provision. Preterm neonatal piglets were nourished exclusively with PN containing either a very high (HG, 30 g/kg/day) or a very low glucose (LG, 2 g/kg/day) level, and systemically challenged with 10^9^ CFU/kg *S. epidermidis*, the dose leading to clinical symptoms but moderate acute mortality (experimental design in Fig. 2A). Although no animals were euthanized preschedule, those provided high amounts of glucose (HG) showed signs of septic shock at 12 h including lethargy, discoloration and tachypnea. Moreover, HG piglets had a quicker passage of meconium compared with animals provided low-glucose (LG), a common physiological stress response in the perinatal period (Fig. 2B). In addition, plasma albumin levels were two times lower in HG relative to LG (P<0.01) to indicate stress-induced changes in liver protein synthesis, vascular permeability or renal dysfunction. This was accompanied by impaired blood bacterial clearance dynamics from 3-12 h in the HG group (Fig. 2C). The effects of *S. epidermidis* infection on blood gases and acidity over time were characterized by decreased pH and acid buffering capacity as well as increased pCO_2_. Importantly, LG reduced blood acidification and respiratory acidosis, relative to HG, and preserved blood acid buffering capacity (Fig. 2D-F). Taken together, restricted glucose supply during neonatal bacterial infection provided acute clinical benefits.

**Figure 2.**
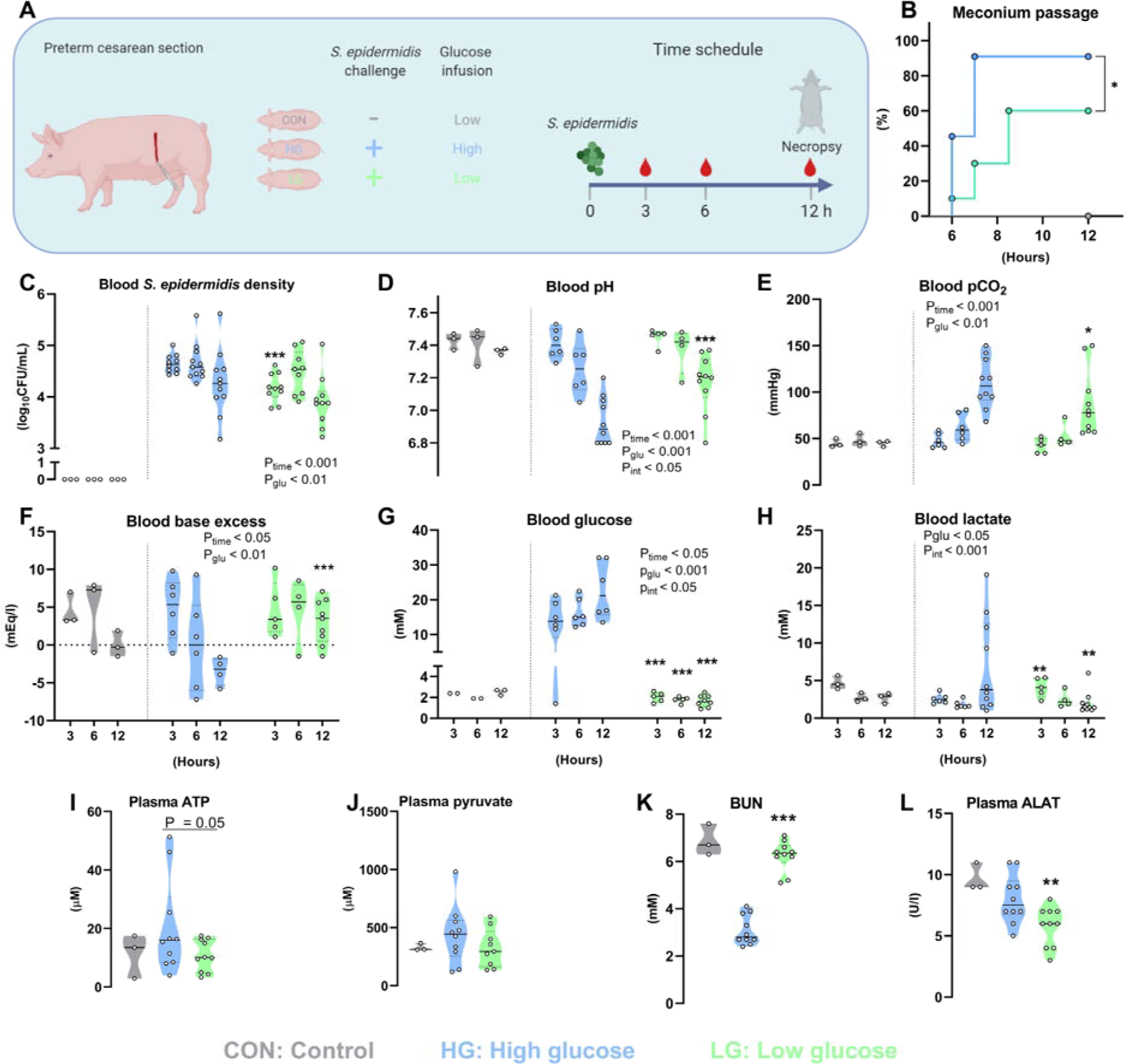
Parenteral glucose restriction protects *S. epidermidis*-infected preterm piglets from sepsis. (**A**) Preterm newborn piglets were nourished exclusively with PN containing high (21%, 30 g/kg/day, HG) or low (1.4%, 2 g/kg/day, LG) glucose concentrations (n = 10-11 per group,) intra-arterially infected with 10^9^ CFU/kg *S. epidermidis*, and cared for 12 h post-infection or until clinical signs of sepsis. Uninfected animals (n = 3) receiving low glucose PN served as a reference and were not included in the statistics. (**B**) Time of first passaged meconium after infection. (**C**) *S. epidermidis* density from blood collected by jugular venous (3-6 h) or heart puncture (12 h), by counting colony-forming units after plating onto tryptic soy agar containing 5% sheep’s blood and incubated for 24 h at 37°C. (**D-H**) Blood gas parameters derived from arterial blood samples collected via the umbilical arterial catheter at 3, 6, and 12 h. (**I-L**). Blood biochemical parameters measured in heparinized plasma from arterial blood collected at 12 h. Data are presented as cumulative hazard curve (B) or violin dot plots including median (solid line) and interquartile range (dotted lines) (C-L). Data are analyzed using a Mantel-Cox test (B) or a linear mixed-effects model (C-L) including an interaction between group and time post-infection (C-H). All analyzed data represents two independent litters. P_time_, P_glu_, and P_int_ denote probability values for effects over time, group effect (HG vs. LG) and interaction effect between time and group in the linear mixed-effects interaction model, respectively. *, **, *** P < 0.05, 0.01, and 0.001, respectively, compared with HG group at the same time point.

Unsurprisingly, HG piglets were hyperglycemic (blood glucose of 10-20 mM) with an increasing trend over time, whereas the LG nourishment paradigm led to hypoglycemia with blood glucose levels around 2 mM and a decreasing time trend (Fig. 2G). We observed a similar pattern for blood lactate (Fig. 2H), where 40% of animals in the HG group had levels above 10 mM, indicating accelerated circulating glycolysis and lactic acidosis, while lactate levels in the LG group decreased over time as it was likely utilized for gluconeogenesis. Despite a large difference in plasma glucose, adenosine triphosphate (ATP) and pyruvate levels only showed minor or no differences between HG and LG groups (Fig. 2I-J). However, blood urea levels were markedly increased in LG relative to HG animals (Fig. 2K), suggesting conversion of exogenous glucogenic amino acids to fuel endogenous glucose production. On the other hand, the plasma activity of alanine aminotransferase, the enzyme responsible for deaminating alanine to pyruvate as an initial step in gluconeogenesis, was decreased in the LG group (Fig. 2L). In summary, high parenteral glucose provision facilitated extensive circulating glycolysis whereas acute metabolic adaptation to exogenous glucose restriction during infection appeared to maintain adequate cellular energy.

During the 12 h course of infection, glucose infusion levels massively interfered with the fate of blood cell subsets. An overall decreasing trend in cell numbers was observed over time for leukocytes, erythrocytes and thrombocytes (Fig. 3A-C), where HG led to a greater loss of total leukocytes and more severe thrombocytopenia. Importantly, HG induced a robust depletion of lymphocytes and neutrophils at 6 h with partial replenishment at 12 h, which was not observed in the LG group (Fig. 3D-E). Monocyte cell numbers tended to be lower at 3 h and replenished at 12h only in the HG group (Fig. 3F). Interestingly, this was associated with distinct temporal changes in the cytokine response to infection. While TNFα, IL10 and IL6 all increased during the course of the infection (Fig. 3G-I), the TNFα and IL6 responses were more pronounced in HG, and conversely IL10 levels increased more over time in the LG group. Collectively, the HG nourishment paradigm induced a more rapid immune response with greater cell loss and evidence of emergency hematopoiesis, prioritizing release of leukocytes but not erythrocytes and thrombocytes from the bone marrow. This may have compromised the regulatory immune response characterized by reduced IL10 secretion.

**Figure 3.**
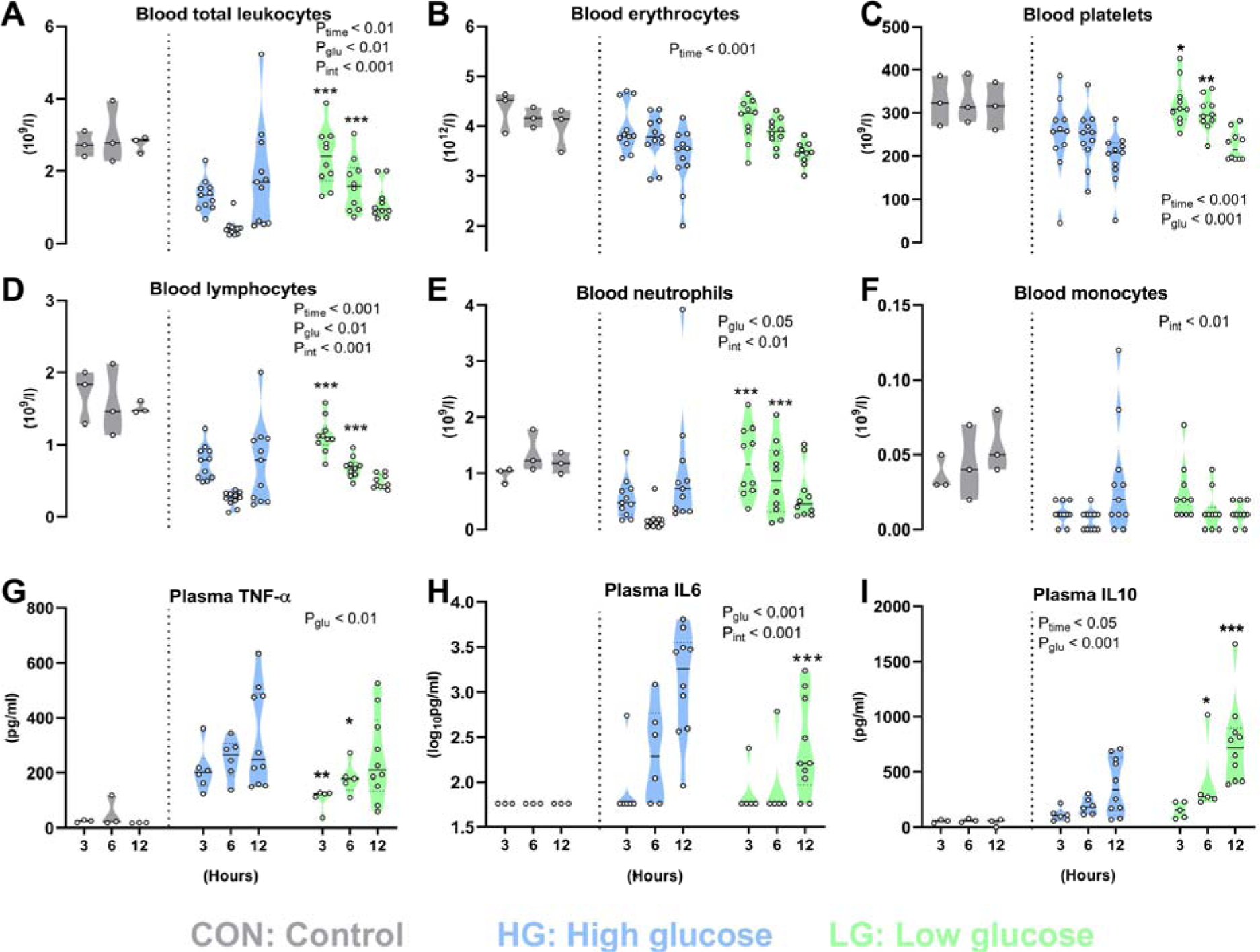
Parenteral glucose restriction protects *S. epidermidis*-infected preterm piglets from excessive inflammation and immune cell loss. (**A-F**) Numbers of hematopoietic cells and major leukocyte subsets in blood samples collected 3-12 h after *S. epidermidis* infusion. (**G-I**). Cytokine levels measured in heparinized plasma from the same blood samples. Data are presented as violin dot plots with median and interquartile range and are analyzed using a linear mixed-effects model including interaction between group and time after infection. All analyzed data represents two independent experiments using separate litters. P_time_, P_glu_, and P_int_ denote probability values for effects over time, group effect (HG vs. LG) and interaction effect between time and group in the linear mixed-effects model, respectively. *, **, *** P < 0.05, 0.01, and 0.001, respectively, compared with HG group at the same time point.

Although glucose restriction has acute clinical benefits with reduced glycolysis, systemic inflammation and clinical signs of sepsis, this practice led to hypoglycemia and may have negative effects on the preterm brain, relying on steady supplies of glucose for proper development. As such, alternative strategies to manipulate the immune-metabolic response to infection in a normo- or hyper-glycemic state should be investigated.

### Glycolysis inhibition decreases inflammatory response to *S. epidermidis in vitro*

Having shown that immune cell metabolism, especially glycolysis, is closely connected to inflammation and clinical fate during neonatal bacterial infection, we aimed to identify a clinically relevant treatment to prevent sepsis and exaggerated aerobic glycolysis beyond glucose restriction. First, we tested the well-known glycolysis inhibitors rapamycin (10 nM, targets mTOR pathway), dichoroacetate (DCA, at 10 mM, targets pyruvate dehydrogenase kinase) and FX11 (100 µM, targets lactate dehydrogenase) for their capacity to reduce inflammatory responses in preterm pig cord blood challenged with *S. epidermidis*. TNFα response was lower when each of the inhibitors was added to cord blood, but DCA seemed to have higher inhibitory potency across the two bacterial doses challenged (Fig. 4A). We proceeded with a dose-finding test for DCA, a short half-life and water-soluble small molecule, widely used for cancer and diabetic patients (29) to suppress inflammation with limited adverse effects (30). At a dose of 10 mM, DCA decreased *S. epidermidis*-induced TNFα secretion more effectively than lower doses (Fig. 4B). DCA at 10 mM but not lower doses tended to be more efficient in decreasing expressions of hexokinase 2 (*HK2*, enzyme facilitating first reaction of glycolysis pathway) and *CXCL8* (pro-inflammatory chemokine), and increasing expression of *IL10* (anti-inflammatory cytokine, Fig. S2A-C). Of note, preterm cord blood incubated with DCA had increased neutrophil phagocytic capacity under both normo- and hyper-glycemic conditions (Fig. 4C). We further performed RNA-seq analysis of *S. epidermidis* stimulated cord blood with or without DCA addition (Fig. 4D-H, Tables S1-8 and Fig. S3) and observed clear patterns of differently expressed genes in control vs. *S. epidermidis-*stimulated samples (90 DEGs) as well as stimulated samples without vs. with DCA (239 DEGs). *S. epidermidis* stimulation up-regulated genes and pathways related to inflammation, innate and adaptive immune activation and down-regulated genes involved in OXPHOS (Fig. 4E-F). Conversely, comparing the two bacteria-stimulated groups, DCA treatment up-regulated anti-inflammatory pathways, pathways related to OXPHOS and mitochondrial ATP synthesis, and down-regulated pathways related to inflammatory responses (Fig. 4E-G). DCA treatment also increased genes related to endocytosis and phagocytosis (Fig. 4H). Collectively, DCA appeared capable of inhibiting infection-induced immune cell glycolysis and inflammation, and was therefore selected as our drug candidate for preventing neonatal sepsis under normo- and hyper-glycemic conditions.

**Figure 4.**
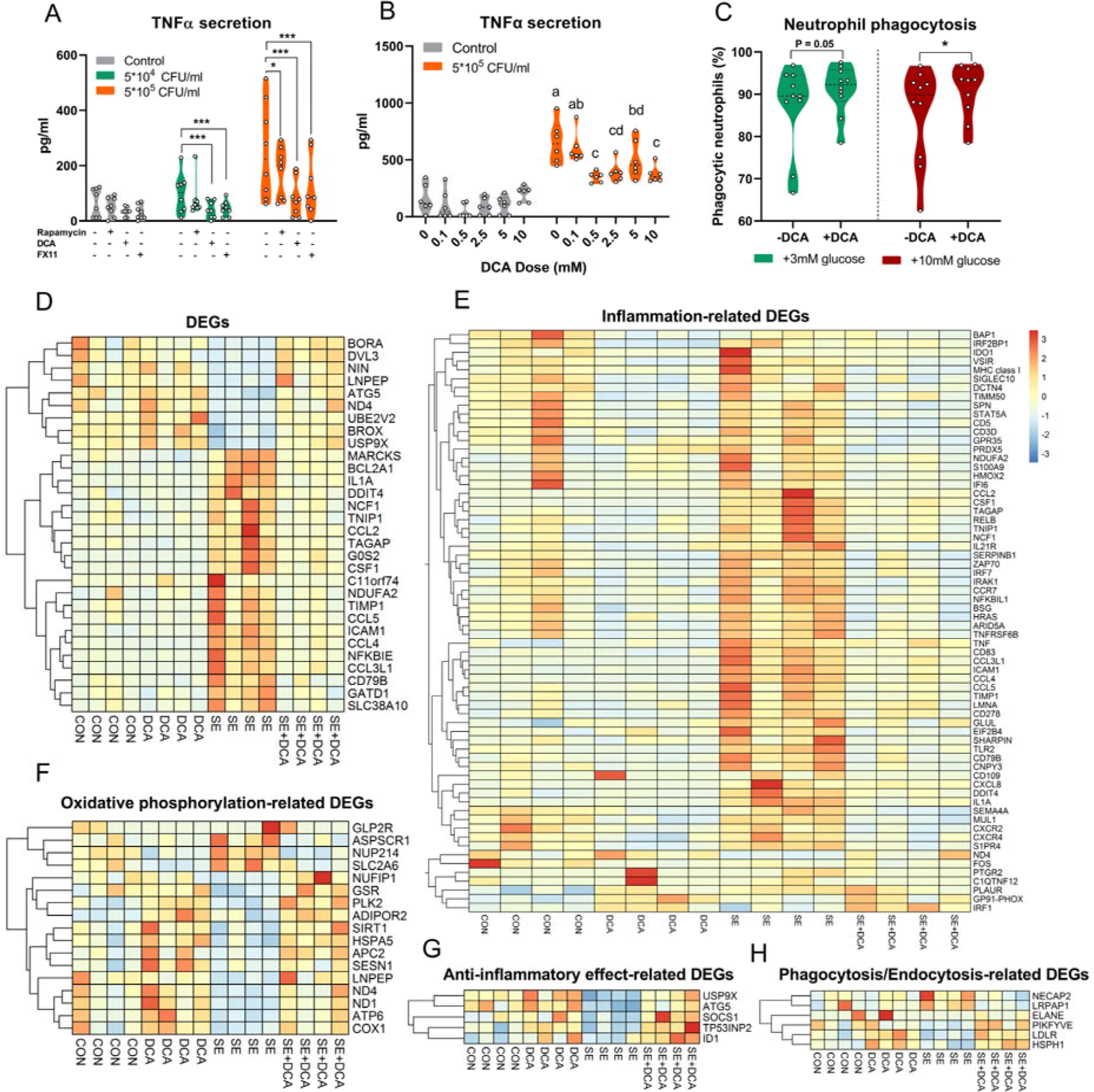
Glycolysis inhibition decreases inflammation in *S. epidermidis*-challenged preterm cord blood. (A-B) TNFα levels in cord blood of preterm piglets (n = 6) following stimulation with *S. epidermidis* (5×10^4^ -5×10^5^ CFU/ml in (A) and the later dose in (B)) with and without presence of glycolysis inhibitor (rapamycin-500 nM, DCA-10 mM in (A) and 0.1-10 mM in (B), FX11-100 µM) at 37°C and 5%CO_2_. incubation (n = 6). (C) Cord blood neutrophil phagocytic capacity (n = 10, from two independent litters) following addition of 10mM DCA or equal volume of sterile water at normo- or hyper-glycemic conditions (added 3 or 10 mM glucose, respectively). *In vitro* phagocytosis assay was performed by incubation samples with pHrodo-conjugated *E.coli* for 30 min at 37°C and 5%CO_2_ and analyzed by flow cytometry. (D-H) Heatmaps from transcriptomic analyses of cord blood samples with/without *S. epidermidis* (5×10^5^ CFU/ml) and DCA incubation (n = 4, from (A) experiment). (D) Top 30 DEGs from the comparison between control vs. *S. epidermidis-*challenged samples. Selective (E) inflammation-, (F) oxidative phosphorylation-, (G) anti-inflammatory effect-, (H) phagocytosis and endocytosis-related DEGs, obtained from the comparison between *S. epidermidis-*stimulated samples without vs. with DCA addition. Normalized expression levels of DEGs were depicted in colors from blue (low) to red (high). Data in (A-C) are presented as violin dot plots with median and interquartile range and analyzed using linear mixed-effect model with inhibitor treatment as a fixed factor and pig ID as the random factor. Transcriptomic data were analyzed by DESeq2 package in R using Benjamini-Hochberg (BH)-adjusted P-value <0.1 as cut-off and further false discovery rate adjustment (FDR, α = 0.1) to convert into q values. *, **, *** P < 0.05, 0.01, and 0.001, respectively, compared with corresponding controls without inhibitor. DCA, dichloroacetate; DEGs: differentially expressed genes; SE*, S. epidermidis*.

### DCA reduces inflammation and improves bacterial clearance during normoglycemia but does not prevent sepsis

Having identified glycolysis inhibition by DCA as a potential alternative to glucose restriction, we again utilized the preterm pig *S. epidermidis* infection model to test the ability of DCA to modulate clinical and molecular outcomes during neonatal infection. The animals were provided with standard (STG, considered current clinical practice, 14.4 g/kg/day) or high parenteral glucose levels (HG, 30 g/kg/day), as well as DCA treatment (50 mg/kg, approximately 10 nM in the circulation, similar to *in vitro* data) or saline control shortly after *S. epidermidis* infusion (experimental design in Fig. 5A). We hypothesized that standard glucose provision as well as DCA treatment would protect against sepsis in the absence of hypoglycemia.

**Figure 5.**
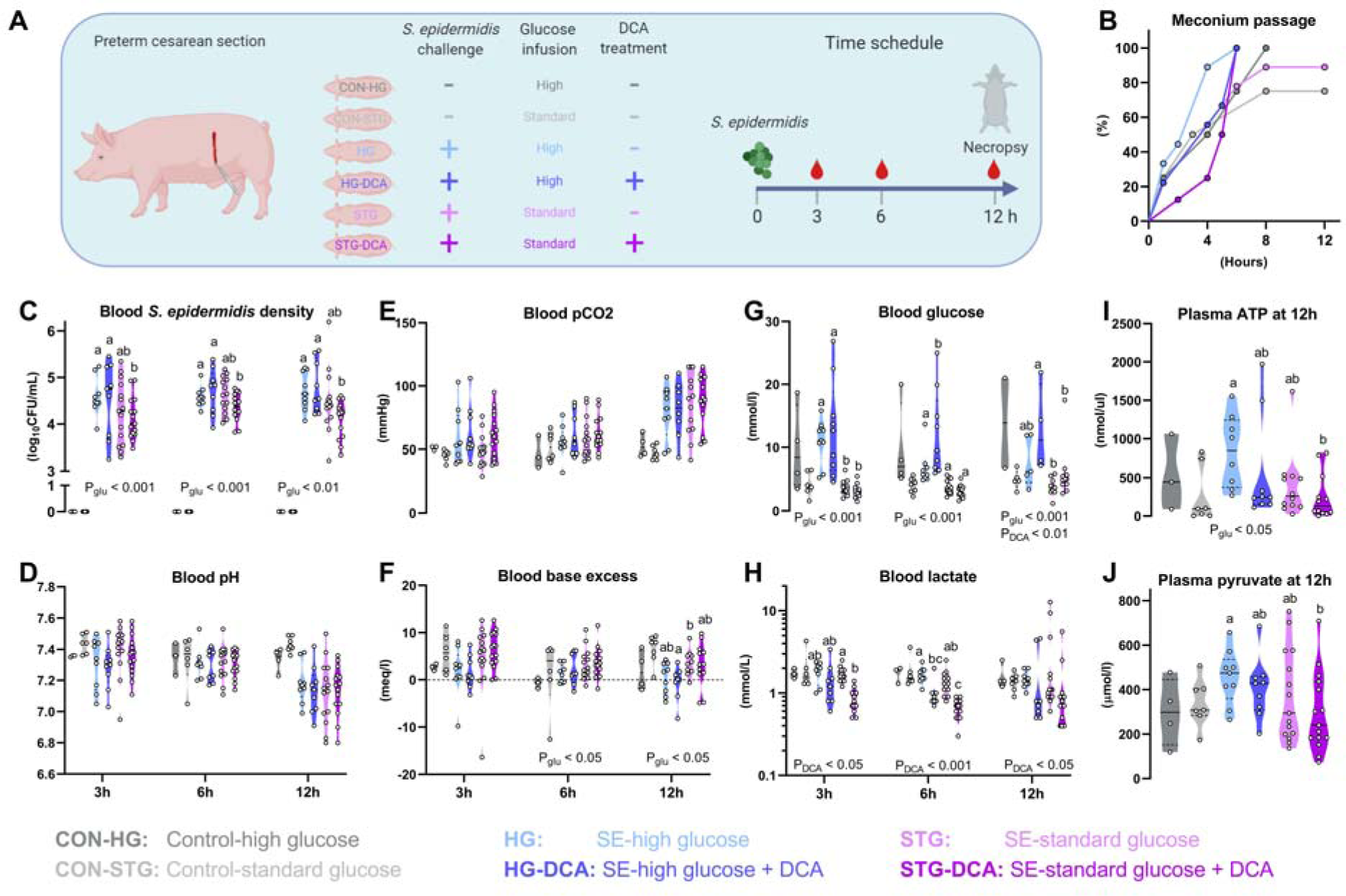
The impact of parenteral glucose levels and glycolysis inhibition by DCA on clinical response to *S. epidermidis* infection. (**A**) Preterm newborn piglets were nourished exclusively with PN containing high (21%, 30 g/kg/day) or standard (10%, 14.4 g/kg/day) glucose concentrations, intra-arterially infected with 10^9^ CFU/kg *S. epidermidis*, followed by saline or DCA treatment (50 mg/kg) 30 min post-infection (n = 9-15/group). Uninfected animals receiving either high or standard PN glucose (n = 4 and 7, respectively) served as reference and were not included in the statistical analysis. (**B**) Time of first passaged meconium after *S. epidermidis* infection. (**C**) *S. epidermidis* density from blood collected by jugular venous (3-6 h) or heart puncture (12 h), by counting colony-forming units after plating onto tryptic soy agar containing 5% sheep’s blood and incubated for 24 h at 37°C. (**D-H**) Blood gas parameters in arterial blood at 3-12 h. (**I-J**). Plasma ATP and pyruvate levels in heparinized plasma from arterial blood at 12 h. Data are presented as cumulative hazard curve and analyzed by Mantel-Cox test (B) or violin dot plots including median and interquartile range and analyzed separately at each blood sampling time point by linear mixed-effect model, including interaction between glucose and DCA (C-J). All analyzed data represent three independent litters. Among infected groups, P_DCA_ and P_glu_ at each time point denote probability values for overall effects of DCA and glucose among the four infected groups in the linear mixed-effect model. Values at each blood sampling time point not sharing the same letters are significantly different (P < 0.05).

The meconium passage time was generally more rapid in this experiment including uninfected controls. Nevertheless, delayed meconium passage was observed in the STG group compared with HG (P<0.05, Fig. 5B) in line with the previous *in vivo* experiment. However, this difference was not present in DCA treated animals. Importantly, bacterial clearance 3-12h post-infection was enhanced in STG groups, relative to HG, while DCA improved bacterial clearance only under STG conditions (Fig. 5C). However, the sepsis indicators blood pH and pCO_2_ were similar across the four infected groups even though there were indications of better blood acid-base balance in STG-DCA pigs across 3-12 h post-infection (Fig. 5D-F). Unsurprisingly, most of the animals in the two infected groups provided high PN glucose were hyperglycemic and most of those in the two infected groups provided the lower standardized PN glucose were normoglycemic. Further, we detected an interesting trend of decreased glucose over time in HG pigs but not in HG-DCA pigs (significantly higher at 6h in HG-DCA pigs, Fig. 5G). This suggested that the HG pigs utilized blood glucose for glycolysis during infection whereas glycolysis inhibition resulted in the consistent hyperglycemic conditions in HG-DCA pigs. Blood lactate was reduced effectively by either lowering PN glucose supply or DCA treatment, reflecting the pyruvate dehydrogenase kinase inhibitory mechanism of action of DCA, with an indication of lowest levels in STG-DCA pigs (Fig. 5H). In parallel, plasma pyruvate and ATP levels, reflecting the degree of energy production enhanced by glycolysis during infection, were reduced in the STG groups, particularly in combination with DCA treatment (Fig. 5I-J). This corroborated the previously presented *in vitro* data.

Reduced PN glucose and DCA interventions also exerted differential effects on blood immune cell subsets and cytokines. In accordance with the previous experiment, infection caused significant reductions of all subsets of immune cells, erythrocytes, recticulocytes and thrombocytes (Fig. 6 A-G). Lowering PN glucose from high to standard levels preserved fractions of neutrophils, lymphocytes, thrombocytes and reticulocytes (Fig. 6 C-E,G). Only STG-DCA treatment showed highest levels of lymphocytes over time, suggesting that DCA only exerts a beneficial effect under normoglycemia. Conversely, the HG-DCA animals had the most severe drops of erythrocytes, thrombocytes and lymphocytes, possibly suggesting the negative impact of more severe hyperglycemic conditions, relative to the HG animals. In line with hematological parameters, plasma levels of IL6, but not IL10, in HG-DCA animals were highest, relative to the remaining three groups (Fig. 6 H-I).

**Figure 6.**
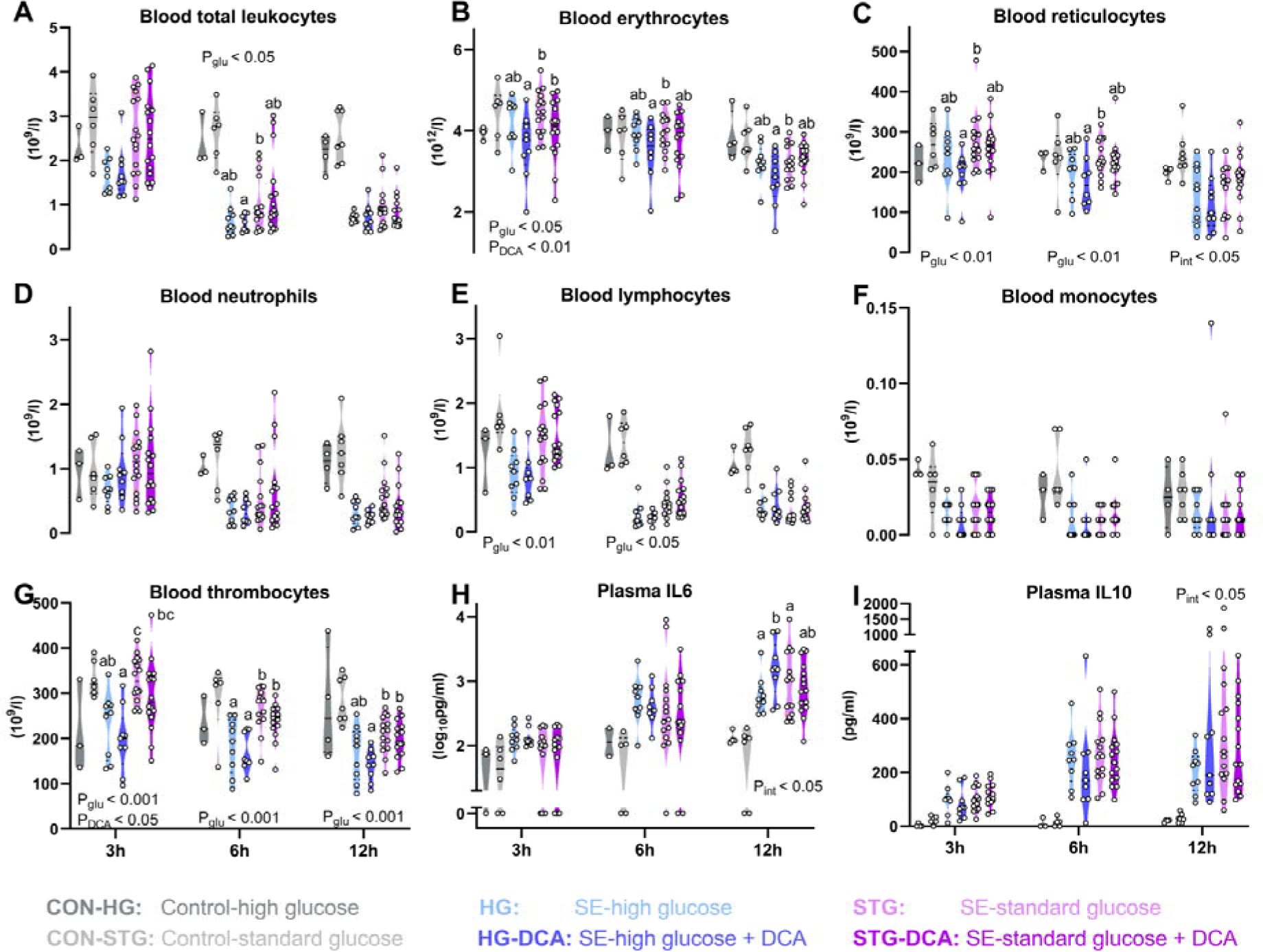
The impact of parenteral glucose levels and glycolysis inhibition by DCA on cellular and cytokine responses to *S. epidermidis* infection. (**A-G**). Numbers of hematopoietic cells and major leukocyte subsets in blood 3, 6, and 12 h after *S. epidermidis* infusion. (**H-I**). Plasma cytokine levels from the same blood samples. Data are presented as violin dot plots with median and interquartile range and are analyzed separately for each blood sampling time poing using a linear mixed-effect model including glucose and DCA interaction. All analyzed data represents three independent experiments using separate litters. Among infected groups, P_DCA_, P_glu_ and P_int_ at each time point denote probability values for overall effects of DCA, glucose and their interaction, respectively, among the four infected groups in the linear mixed-effect model. Values at each blood sampling time point not sharing the same letters are significantly different (P < 0.05).

To better understand the effects of standard glucose supply and DCA at molecular levels, a subset of blood samples at 12 h post-infection were used for whole-transcriptome analyses (Fig. 7 and Tables S9-17). *S. epidermidis* infection induced dramatic blood transcriptome changes (21.2% of annotated genes), with 1967 down- and 2011 up-regulated DEGs, leading to elevated pathways related to innate immunity (TLR, NOD signaling) and early phase of Th1 polarization (chemokine and TNF signaling) and down-regulated adaptive immune pathways (T and B cell receptor signaling, Fig.7A-B, Table S15-17). Multiple metabolism-related genes/pathways were also down-regulated by infection, including fatty acid degradation (Fig.7B). Comparisons among the four infected groups (Table S9-14) showed that HG pigs possessed a distinct profile of inflammation-related genes with half of the DEGs being highly up-regulated and the other half being down-regulated, relative to the remaining three groups (Fig. 7C). Surprisingly, HG-DCA pigs with the worst clinical outcomes possessed a similar blood transcriptome profile to STG and STG-DCA pigs. HG pigs had increased levels of multiple genes related to energy metabolism and ATP synthesis, when compared to STG (Fig. 7D) or HG-DCA pigs (Fig. 7E). These data suggest that high circulating glucose levels accelerated metabolic pathways to synthesize ATP fueling excessive inflammatory responses to infection. Conversely, reducing PN glucose intake from high to standard levels or using DCA treatment conveyed similar changes at transcription level to the direction of less inflammation and energy metabolism. These in combination with other data imply that the detrimental impact of DCA during high PN glucose supply on clinical outcomes were likely derived from the more severe hyperglycemia induced by the inhibitory effects of DCA on blood cellular glucose uptake.

**Figure 7.**
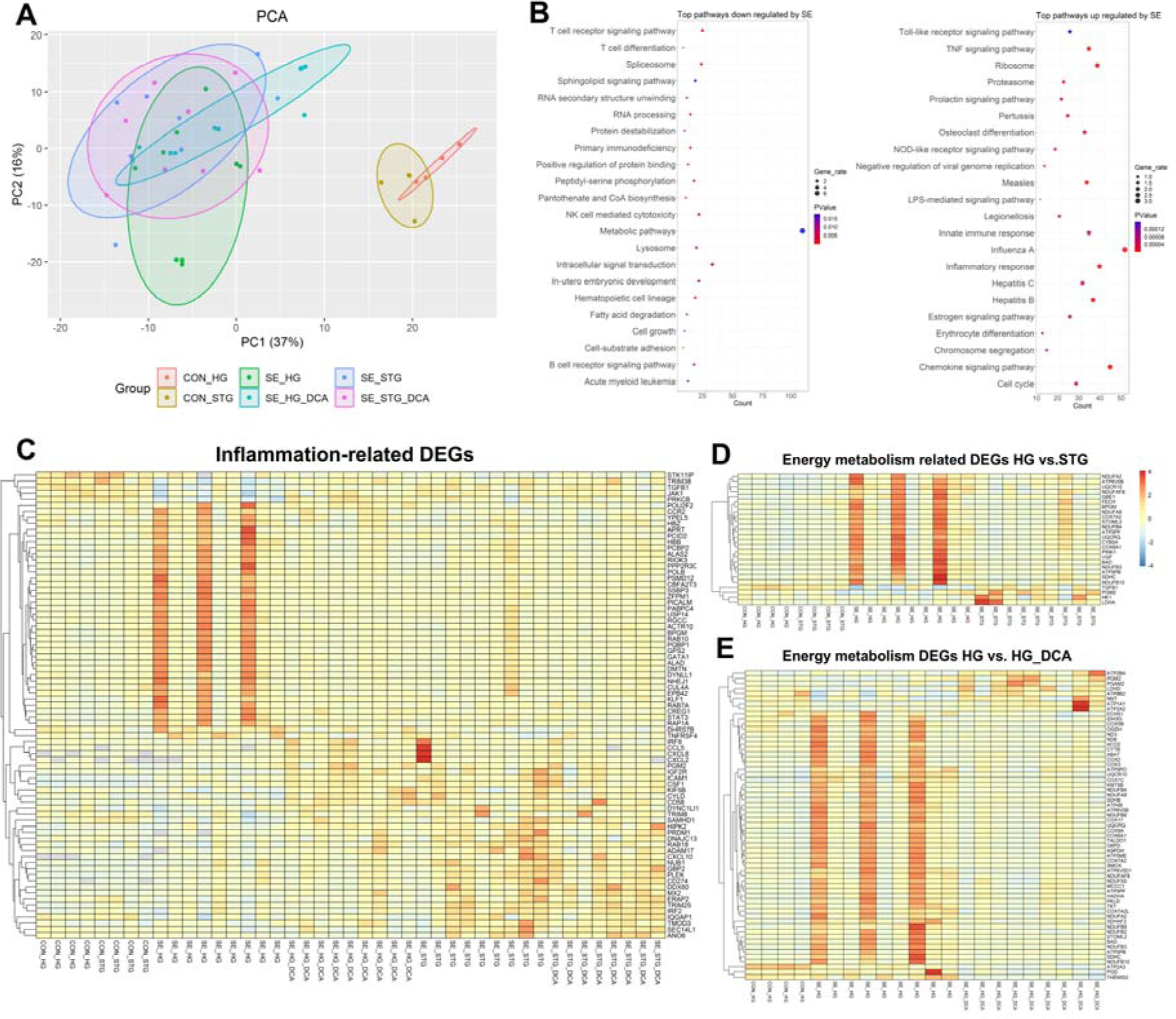
Blood transcriptomic responses to *S. epidermidis* infection and the effects of parenteral glucose levels and glycolysis inhibition by DCA. **(A)** Principal component analysis of the blood transcriptome at 12h in control or infected pigs nourished with high or standard parenteral glucose regime, with/without DCA treatment. **(B)** Top pathways regulated by *S. epidermidis* infection analyzed by KEGG and GO pathway enrichment analysis using DAVID database, and DEG counts displayed in X axis. **(C-D)** Heatmaps including inflammation and energy metabolism-related DEGs between HG vs. STG animals. **(E)**. Heatmap incuding energy metabolism-related DEGs between HG vs. HG-DCA animals. Analyses included 4 animals per control group and 8-9 per each infected group in two independent litters. Normalized expression levels of DEGs were depicted in colors from blue (low) to red (high). All the statistics was performed by DESeq2 with FDR adjusted by Benjamini-Hochberg (BH) procedure using α = 0.1 as the threshold.

In summary, whereas standard relative to high glucose provision reduced the acute stress response to *S. epidermidis* infection and improved bacterial clearance from the blood, it failed to provide the same protection against sepsis as bona fide glucose restriction. Moreover, DCA treatment appeared to offer further benefits in infected animals on standard glucose provision, while it exacerbated the hyperglycemic condition and inflammation during high glucose provision.

## Discussion

A delicate balance of metabolic and immune responses determines how neonates manage to survive serious infections (8). Here, we identified bacterial dose-dependent immunometabolic thresholds *in vitro* and *in vivo*, and uncovered the modulatory roles of systemic glucose provision and cellular energy metabolism in regulating inflammation and sepsis outcomes in a clinically-relevant animal model of neonatal bloodstream infection. First, the metabolic responses of cord blood and preterm experimental animals to *S. epidermidis* were dose-dependent, whereby glycolysis and OXPHOS markers increased and decreased, respectively, with increasing bacterial dose only until a certain dose, after which they were normalized to levels in controls. This is a demonstration of the Warburg effect taking place in activated immune cells (31) and defines the immunometabolic thresholds from immune tolerance to activation and later immunoparalysis coupled with perturbed glucose homeostasis. Second, a proof-of-concept study showed that high parenteral glucose provision in infected individuals was clearly detrimental, via induced hyperglycemia, accelerated glycolysis producing lactate and ATP, fueling inflammatory responses and leading to sepsis. In contrast, parenteral glucose restriction caused hypoglycemia but reduced inflammation and protected against sepsis. Third, cord blood stimulated with *S. epidermidis* and glycolysis inhibitors showed clear effects of glycolytic inhibition to reduce inflammation, and to enhance OXPHOS and neutrophil phagocytosis. Finally, we found that effects of glycolysis inhibition by the pyruvate dehydrogenase kinase inhibitor DCA in infected individuals depended on the level of glucose provision. DCA increased inflammation and prompted more severe hyperglycemia during high glucose provision, whereas it decreased inflammation and improved bacterial clearance during standard glucose provision, albeit without preventing sepsis. The mechanistic insights from the current study suggest that parenteral glucose restriction could be a lifesaving therapy for infected preterm infants despite causing temporary hypoglycemia.

Increasing bacterial dose *in vitro* and *in vivo* led to the switch from tolerant to resistant responses (increased cellular glucose uptake producing lactate, decreased OXPHOS-related genes and Treg levels), and finally to immunoparalysis or death. The tolerant status in the current study is similar to the impaired immunometabolic responses to LPS or bacteria of preterm vs. term monocytes (10, 32), or that of preterm infants with vs. without sepsis (4). Increased bacterial dose beyond the resistance threshold decreased the ratio of pro- vs. anti-inflammatory genes, normalized Treg levels and OXPHOS-related genes to that in controls (immunoparalysis), which likely occurred when blood cells used up all their energy stores. This is similar to the immunosuppression in late stages of sepsis in infants and elderly (14, 33), which predisposes to secondary infections. To further test effects of glucose provision and pharmacological glycolysis inhibition, we selected the *S. epidermidis* dose exerting resistant responses and sepsis signs without significant acute mortality. Relative to infected animals with restricted parenteral glucose, those with high parenteral glucose supply had hyperglycemia, thrombocytopenia, leukopenia, lower blood pH, impaired bacterial clearance, and higher levels of blood lactate, pCO2, ATP, and inflammatory cytokines. Hyperglycemia in infected adult animals is known to impair monocyte chemotaxis and neutrophil phagocytosis, thereby decreasing systemic bacterial clearance and increasing sepsis risk (34, 35). The mechanisms for this is unclear, despite few studies showing hyperglycemia-induced impaired IgG influx and complement protein release, which are needed for opsonization (36). Further, hyperglycemia likely enhanced glucose uptake to accelerate glycolytic activity, in turn increasing lactate and ATP production used for inflammatory responses. The combination of elevated glycolysis and hyperglycemia-induced impaired phagocytosis may explain poorer sepsis outcomes in these infected animals.

In contrast, restricted parenteral glucose caused hypoglycemia but completely prevented infected animals from elevated glycolysis, excessive inflammation and sepsis. This solution may not be practical for hospitalized infants due to the fear of hypoglycemia-induced brain injury (17). Therefore, we postulated that any other ways of reducing glycolysis may be beneficial for both infection and sepsis outcomes without causing hypoglycemia. This was in principle challenging as phagocytes are also dependent on glycolysis to clear bacteria via phagocytosis (37-39). Via screening various drugs, we identified DCA, a pyruvate dehydrogenase kinase inhibitor, with short half-life that enhances OXPHOS and decreases glycolysis, thereby potentially reducing ATP production and inflammation during infection (40). It has also been used for adult cancer patients (41). We found that DCA was detrimental in hyperglycemic but moderately beneficial in normoglycemic infected animals. Specifically, on high PN glucose background, DCA induced more severe hyperglycemia, higher levels of inflammatory cytokines and reduced bacterial clearance, suggesting its overall detrimental effects likely derived from the decreased cellular glucose uptake during hyperglycemia. In contrast, reducing systemic glucose provision more moderately reduced dysglycemia, glycolysis, lactic acidosis and inflammation via restricted cellular glucose influx, while DCA use on this lower parenteral glucose background further dampened inflammation via its inhibitory effects on glycolysis and lactate production. Importantly, despite evoking temporary glycolysis inhibition, DCA treatment during normoglycemia also enhanced *in vivo* bacterial clearance, likely via the enhanced neutrophil phagocytosis, as evidenced by transcriptomic responses and phagocytosis test in DCA-treated cord blood and also previous studies (42, 43). Clearly, the interaction between parenteral glucose supply and DCA mechanistic actions determined the inflammatory outcomes, suggesting careful blood glucose monitoring during DCA use.

Importantly, whether with or without additional DCA treatment, standardization of PN glucose supply to that recommended in clinical neonatal guidelines (17) could not prevent infected animals from showing clinical signs of sepsis, including acidemia (decreased blood pH) and respiratory acidosis (increased pCO2) despite ameliorated inflammatory effects. Our results thus challenge the appropriateness of this international guideline, as this does not consider detailed PN glucose regimes during neonatal infection. Hence, there is a great need for clinical trials testing lower PN glucose provisions in infected preterm infants. If these lower glucose interventions cannot further decrease sepsis risk, restricted parenteral glucose during neonatal infection as shown in our current study may be a solution to prevent life-threatening sepsis despite causing temporary hypoglycemia.

## Materials and Methods

### *S. epidermidis* culture preparation

*S. epidermidis* (WT-1457) was prepared from frozen stock. Bacteria were previously cultured in heart infusion broth (HIB). Bacterial concentration and optical density (OD) in the stock was pre-determined by CFU counting following bacterial stock washing in PBS (Sigma-Aldrich), resuspension and dilution in PBS at 4°C, and 20µl spotting in triplicates on a blood agar plates for overnight incubation at 37°C. Right before *in vitro* experiments, bacterial stock was thawed and diluted in PBS at 4°C to reach the desired concentrations for cord blood stimulation, based on the pre-determined concentration in the stock. For *in vivo* experiments, 30 ml tryptic soy broth was inoculated with 500 µl *S. epidermidis* stock and incubated for 17 h at 37°C and 200 rpm. Culture OD was then measured by spectrophotometry and bacterial concentration estimated based on a previously established OD-to-CFU conversion factor. The culture was centrifuged for 20 min at 3000 × g and bacterial pellet suspended in sterile physiological saline at 3 × 10^8^ CFU/ml. The *S. epidermidis* culture was serially diluted and plated onto tryptic soy agar and incubated overnight at 37°C to verify the actual concentration for each *in vitro* and *in vivo* experiment.

### *In vitro* cord blood stimulation with *S. epidermidis*

Cord blood collected at preterm pig delivery (day 106 of gestation, term at day 117±2) was aliquoted into a sterile 96 well plate (200µl/well) and stimulated with an increasing dose of *S. epidermidis* (5×10^1^-5×10^7^ cells/ml blood) at 37°C with 5% CO_2_ for 2 h. In some experiments, blood was pre-incubated with various concentrations of glycolysis inhibitors (rapamycin, DCA and FX11, all from Sigma-Aldrich, Copenhagen, Denmark). After stimulation, 90 µl blood was stabilized with 200 µl mixture of lysis/binding solution concentrate and isopropanol (MagMax 96 blood RNA isolation kit, Thermofisher, Roskilde, Denmark), and stored at -80°C for later RNA extraction. The remaining blood was centrifuged (2000×g, 10 min, 4°C), and plasma analyzed for cytokines and metabolic targets. In some experiments, cord blood after stimulation was used for flow cytometry analysis of regulatory T cells (Treg). All *in vitro* experiments were performed using cord blood from at least 4 preterm animals.

### *In vivo S. epidermidis* infection in preterm pigs

The animal studies and experimental procedures were approved by the Danish Animal Experiments Inspectorate (license no. 2020-15-0201-00520), which complies with the EU Directive 2010/63. All piglets (cross-bred, Landrace x Yorkshire x Duroc) were delivered by elective cesarean section at gestational day 106 corresponding to ∼90% of the total length of gestation. Sow’s anesthesia and surgical procedures are described in details elsewhere (44). After delivery, the animals were single-housed in ventilated, heated (37°C) incubators with oxygen supply (1 l/min). For resuscitation, animals received Doxapram and Flumazenil (0.1 ml/kg each drug, intramuscularly), and positive airway pressure ventilation until breathing stabilized. Once stabilized, a 4 Fr gauge catheter was inserted into one of the umbilical arteries under aseptic conditions and fixed at the level of the descending aorta for provision of parenteral nutrition (PN), *S. epidermidis* inoculation and blood sampling. Successfully resuscitated animals were stratified by sex and birth weight, and allocated into treatment groups using random number generation. In all animal experiments, *S. epidermidis* was administered intra-arterially as a 3 min continuous infusion (3.33 ml/kg) within 4 h after birth using a precision infusion pump. The animals were nourished parenterally with Kabiven infusion formula (Fresenius-Kabi, Bad Homburg, Germany) using different glucose concentrations at infusion rates of 6 ml/kg/h. Animals were permanently monitored by experienced caretakers for the duration of the experiments (12-24 h) and euthanized preschedule if presenting with clinical signs of septic shock (e.g. extreme lethargy, discoloration, hypo-perfusion). Blood was collected by jugular venous or heart puncture on sterilized skin for bacteriology and through the umbilical catheter for the remaining analytical endpoints. Scheduled euthanasia was preceded by induction of deep anesthesia and executed by a lethal dose of intra-cardiac barbiturate. Animal caretakers were not blinded to the respective treatment groups, but all endpoint and data analyses (except meconium passage time) were conducted in a blinded fashion.

In the initial *in vivo* bacterial dose-response experiment that served to establish a clinical and metabolic phenotype, animals were randomly allocated to receive saline (CON, n = 13), 10^6^ (n = 7), 10^8^ (n = 14), 10^9^ (n = 10), or 5 × 10^9^ (n = 13) CFU/kg *S. epidermidis*. Animals were nourished with PN containing a standard glucose concentration (10%), corresponding to a daily glucose intake of 14.4 g/kg, and monitored for 24 h including blood collection at 12 and 24 h.

In the subsequent experiment addressing the hypothesis that glucose restriction protected against sepsis, animals were randomly allocated to receive 10^9^ CFU/kg *S. epidermidis* and PN formula containing either a low glucose (LG, 1.4% or 2 g/kg/d, n = 10) or high glucose (HG, 21% or 30 g/kg/d, n = 11) concentration. A third group of reduced sample size served as uninfected controls (CON, n = 3) and received low glucose (1.4%) parenteral formulation. All animals were monitored for 12 h including blood collection at 3, 6, and 12 h.

The final experiment addressed the hypothesis that reducing PN glucose intake from high to standard regimes with or without indirect glycolysis inhibition via DCA administration (directly inhibiting pyruvate dehydrogenase kinase 1, PDK1) would protect against sepsis. Animals were randomly allocated to receive 10^9^ CFU/kg *S. epidermidis* and PN containing standard glucose concentration (STD, 10%, 14.4 g/kg/d, n = 15) without or with DCA (STD-DCA, n = 15), or PN with high glucose concentration (HG, 21%, n = 9) without or with DCA (HG-DCA, n = 9). DCA groups received 50 mg/kg (1 ml solution per kg) DCA intra-arterially exactly 30 min after *S. epidermidis* infusion, whereas DCA controls received an equivalent volume of sterile physiological saline. Some infected STD-DCA animals (n = 7) also received additional DCA (50 mg/kg) at 3 and 6 h post-infection but showed no additional effects relative to those with single DCA treatment, and were therefore pooled to form the STD-DCA group. Besides, animals were randomly allocated to two uninfected control groups receiving either standard (CON-STD, n = 7) or high PN glucose (CON-HG, n = 4). The animals were monitored for 12 h including blood sampling at 3, 6, and 12 h.

### Treg and neutrophil phagocytosis

In an *in vitro* experiment, the frequency of Treg cells in stimulated blood was analyzed as previously described(28). In brief, blood after bacterial stimulation was lyzed to remove erythrocytes, washed with PBS, permeabilized (permabilization buffer, Thermofisher), blocked with porcine serum (Thermofisher Scientific), and stained with a mixture of FITC-conjugated mouse-IgG2b anti-porcine CD4 antibody (clone MIL17), APC-conjugated rat-IgG2a anti-porcine Foxp3 antibody (clone FJK-16s), and analyzed by a BD Accuri C6 flow cytometer (BD Biosciences, USA). Treg was defined as CD4^+^Foxp3^+^ lymphocytes. In another experiment, cord blood pre-incubated with a glycolysis inhibitor DCA under different glycemic conditions was assessed for its phagocytosis capacity as previously described(45). In brief, 100 µl cord blood was stimulated with pHrodo red-conjugated *E.coli* bioparticles (Phagocytosis kit, Thermofisher) at 37°C for 30 min, followed by flow cytometry analysis as mentioned above. Percentage of neutrophils having phagocytic capacity in total number of neutrophils was evaluated.

### Gene expression analysis by qPCR

Total whole blood RNA from *in vitro* and *in vivo* experiments was extracted using MagMAX 96 Blood RNA Isolation Kit (Thermofisher). RNA was then converted to cDNA with the High capacity cDNA reverse transcription kit (Applied Biosystems, USA). Transcription of selected genes related to inflammation, innate and adaptive immunity and energy metabolisms were determined by quantitative polymerase chain reaction (qPCR) using QuantiTect SYBR Green PCR Kit (Qiagen, Netherlands) on the LightCycler 480 system (Roche, Switzerland) with predesigned primers (sequences in Table S18). Primers were designed with Genes database and Primer-BLAST software (National Center for Biotechnology Information, USA). Relative expression of target genes was calculated by double delta Ct method with HPRT1 served as the housekeeping gene.

### Whole transcriptome shotgun sequencing

Whole blood RNA of selected samples from *in vitro and in vivo* experiments was analyzed by whole transcriptome shotgun sequencing, as previously described(28), to profile immunometabolic pathways affected by relevant interventions. Briefly, RNA-seq libraries were constructed using 1000 ng RNA and VAHTS mRNA-seq V3 Library Prep Kit for Illumina (Vazyme, China). The libraries were sequenced on the Illumina Hiseq X Ten platform (Illumina, USA) to generate 150 bp paired-end reads. Quality and adapter trimming of raw reads were performed using TrimGalore (Babraham Binoinformatics, UK). The remaining clean reads (∼ 26 M per sample) were aligned to the porcine genome (Sscrofa11.1) using Tophat2(46). The annotated gene information of porcine genome was obtained from Ensembl (release 99). The script htseq-count(47) was used to generate gene count matrix, followed by analyses of differentially expressed genes (DEGs) using DESeq2(48).

### Plasma cytokines and metabolic targets

Plasma from *in vitro* and *in vivo* experiments were analyzed for porcine specific cytokines using enzyme-linked immunosorbent assay (ELISA, TNFα (DY690B), IL10 (DY693B) and IL6 (DY686, porcine DuoSet, R&D systems, Abingdon, UK), and targets related to energy metabolism. Glucose and lactate were measured by Lactate Assay Kit and Glucose Assay Kit, respectively (all from Nordic BioSite, Denmark). Extracellular ATP and pyruvate levels were measured by the ATP Colorimetric/Fluorometric Assay Kit and the Pyruvate Assay Kit (SigmaAldrich).

### Statistics

All continuous data were analysed using in R studio 3.4.1 (R Studio, Boston, MA). *In vitro* data were analysed by a linear mixed-effect model with treatment as a fixed factor and pig ID as a random factor, followed by Tukey Post-hoc pair-wise comparisons. Survival curves (meconium passages or survival) were analyzed using Matel-Cox log-rank tests. To compare HG vs. LG infected animals, each parameter was fitted in to a linear mixed-effect model with glucose level, time and their interaction as fixed factors and litter and pig ID as random factors, using lme4 and multcomp packages(49). Group comparisons at each time point were also performed with similar models without contributing factors of time of blood sampling and pig ID. For the experiment identifying glucose and DCA effects in infected animals, each parameter at each blood sampling time point was fitted into a linear mixed-effect model with glucose, DCA, and their interaction as fixed factors and pig ID as a random factor. For pair-wise comparisons, Tukey Post-hoc test was used after a linear mixed effect model was applied with treatment as a fixed factor and litter as random factor. An adjusted P-value < 0.05 was regarded as statistically significant. Data are presented as violin dot plots with median and interquartile range. All reported measures were evaluated for normal distribution, and logarithmic transformation was performed if necessary. For transcriptomics, significant DEGs among groups were identified by DESeq2 using Benjamini-Hochberg (BH)-adjusted P-value <0.1 as cut-off. To control type I error, p values tests were further adjusted by false discovery rate (FDR, α = 0.1) into q values(50). Gene ontology and KEGG pathway enrichment analyses were performed using DAVID (51) and a BH-adjusted P-value <0.05 was considered statistically significant. Lists of genes with mean expression levels and adjusted P-values as well as enriched pathways with associated DEGs were listed for each comparison from *in vitro* (Table S1-8), and *in vivo* experiments (Table S9-17). Heatmaps were generated using R package pheatmap.

## Acknowledgements

The authors thank Per Sangild, Thomas Thymann, Ole Bæk, Rene L. Shen, Xiaoyu Pan, Britta Karlsson and Jane C. Povlsen for the assistance in animal experiments and omic data acquirement.

## Author contributions

DNN designed the study. TK, AB, NLH, KAS and DNN performed the animal experiments and laboratory analyses. TM, AB and DNN conducted bioinformatics, statistical analysis and data interpretation. TM and AB managed raw data and generated all figures and tables. TM, AB and DNN drafted the manuscript. All authors contributed to data interpretation, manuscript revision and approval of the final manuscript version.

