## Supplementary Figures for "Parenteral glucose supply and pharmacological glycolysis inhibition determine the clinical fate of infected preterm newborns"

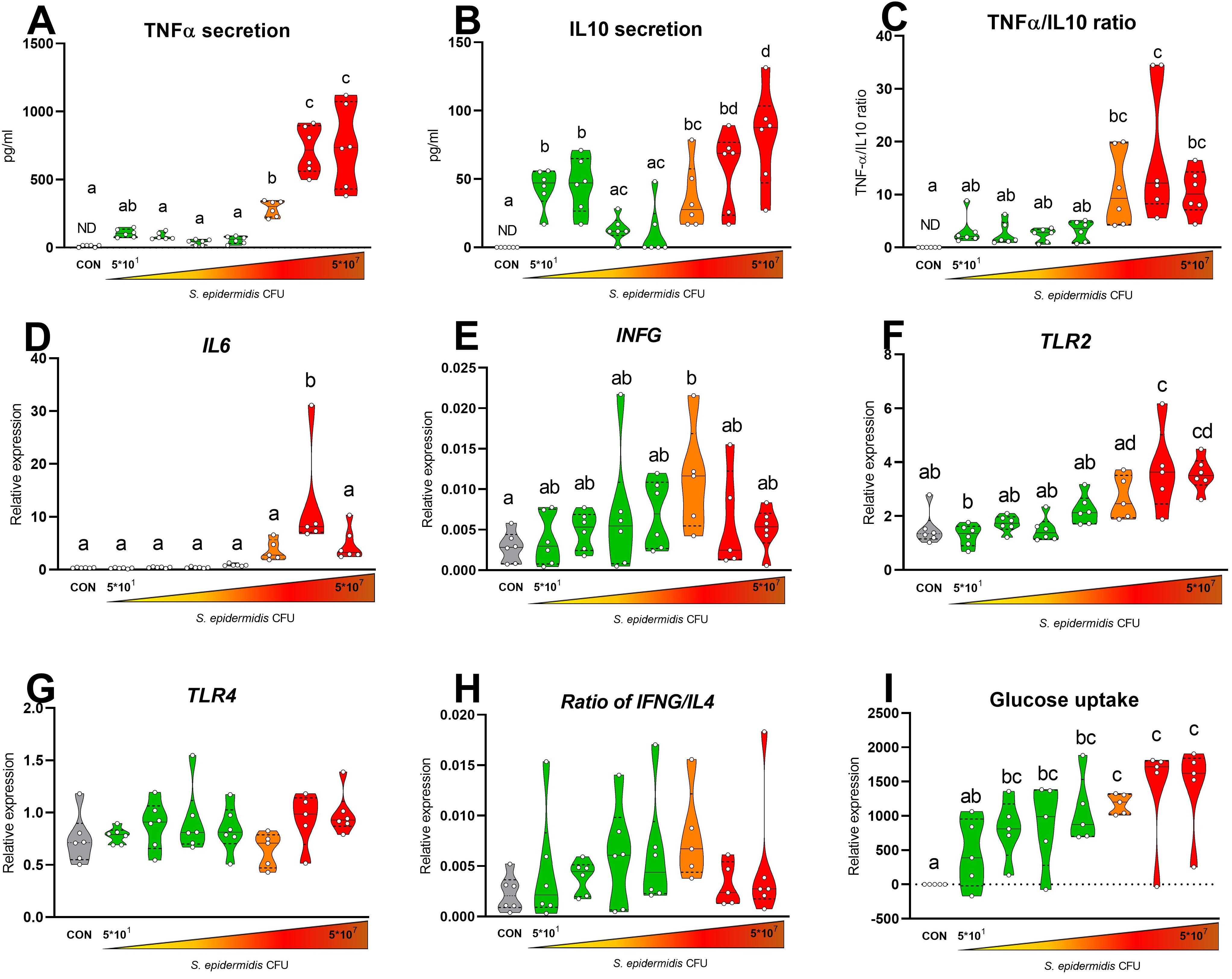

**Figure S1: *In vitro* immunometabolic response of preterm pig cord blood to increasing doses of *S.epidermidis*.** Blood samples were stimulated with bacteria (5×10^1^ -5×10^7^ CFU/mL) for 2 h at 37°C and 5%CO_2_. **(A-C)** Plasma cytokine levels of TNFA, IL10, and TNFA/IL10 ratio (n = 5-6). **(D-H)** mRNA levels of *IL6*, *INFG, TLR2, TLR4,* and *TNGF/IL4* ratio (n = 5-6). (**I**) Cellular glucose uptake measured by the differences in supernatant glucose levels of stimulated and unstimulated samples (n = 5). Data are presented as violin dot plots with median (solid line) and interquartile range (dotted lines) and were analyzed using linear mixed-effect model followed by Tukey Post-hoc comparisons. Values not sharing the same letters are significantly different (P < 0.05).

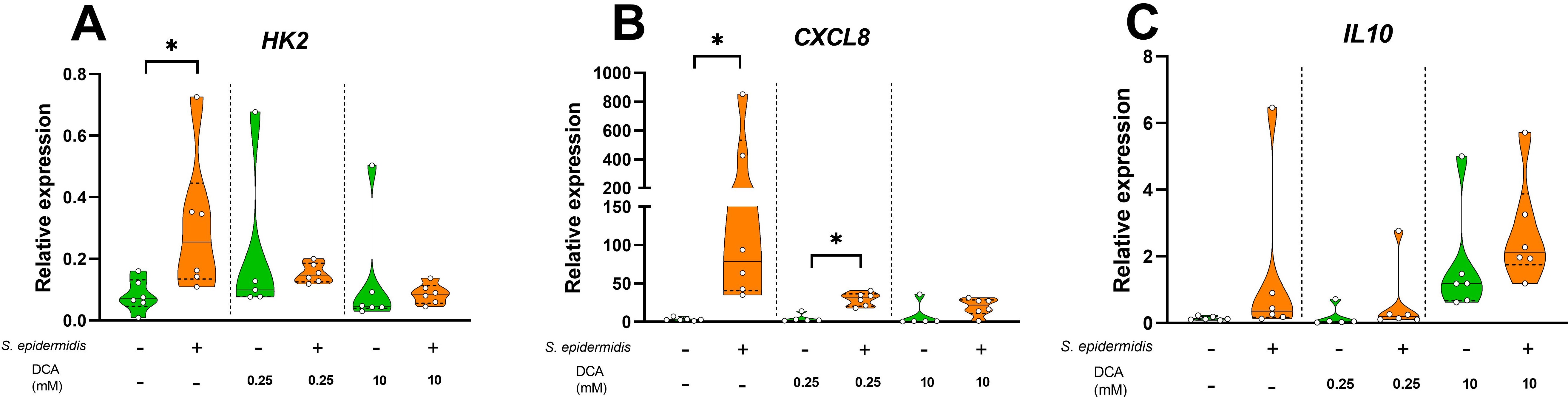

**Figure S2: *In vitro* immunometabolic response of preterm pig cord blood to *S.epidermidis* and dichloroacetate (DCA) supplementation. (A-C)** mRNA levels of *HK2,CXCL8, IL10* of cord blood from preterm piglets stimulated with and without *S. epidermidis* (5×10^5^ CFU/ml), with and without presence of glycolysis inhibitor DCA (0.25mM or 10mM DCA), for 2 hours at 37°C and 5% CO_2_ (n = 5-6). Data are presented as violin dot plots with median (solid line) and interquartile range (dotted lines) and were analyzed using linear mixed-effect model followed by Tukey Post-hoc comparisons. * , P < 0.05.

**A**

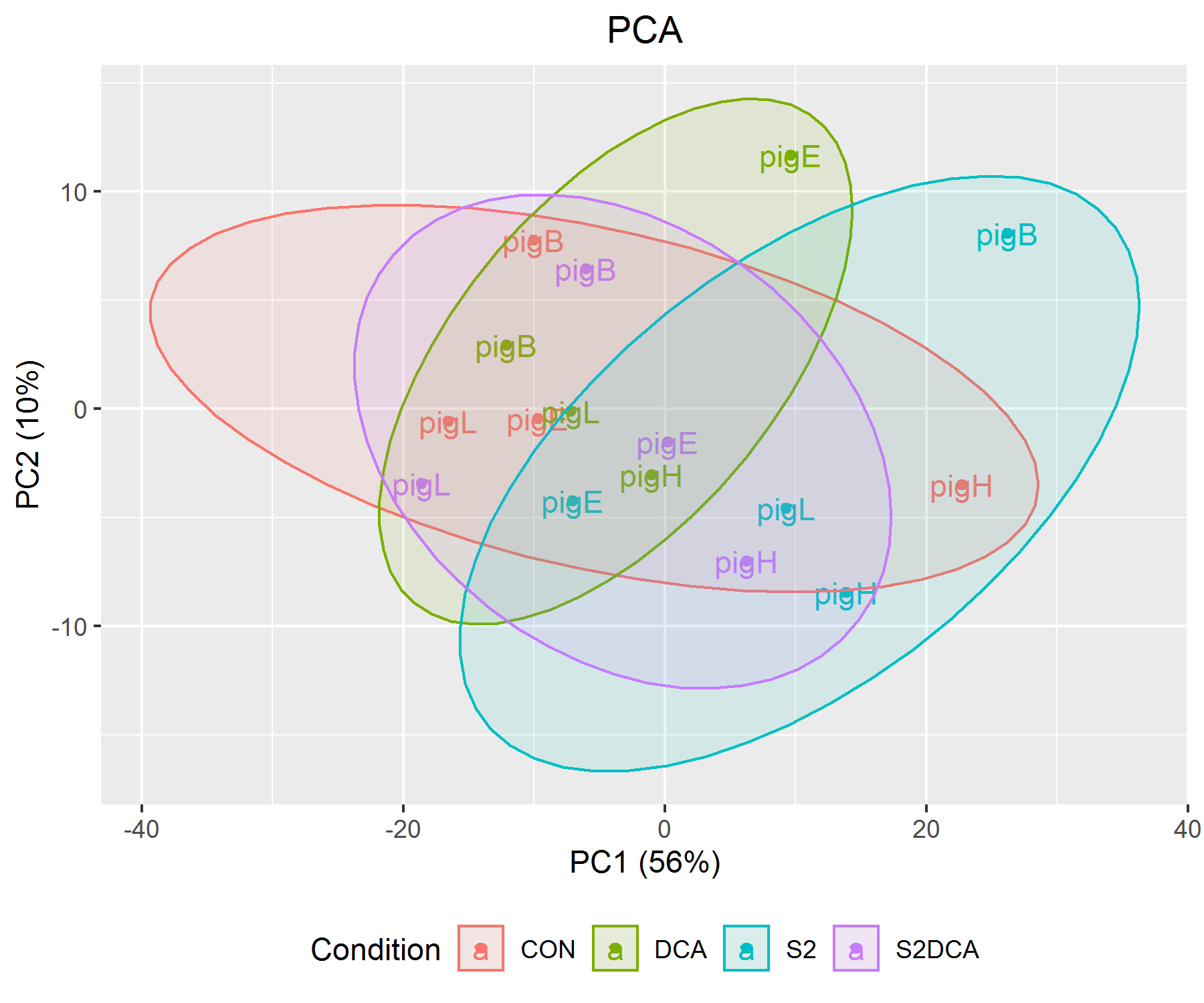

**B**

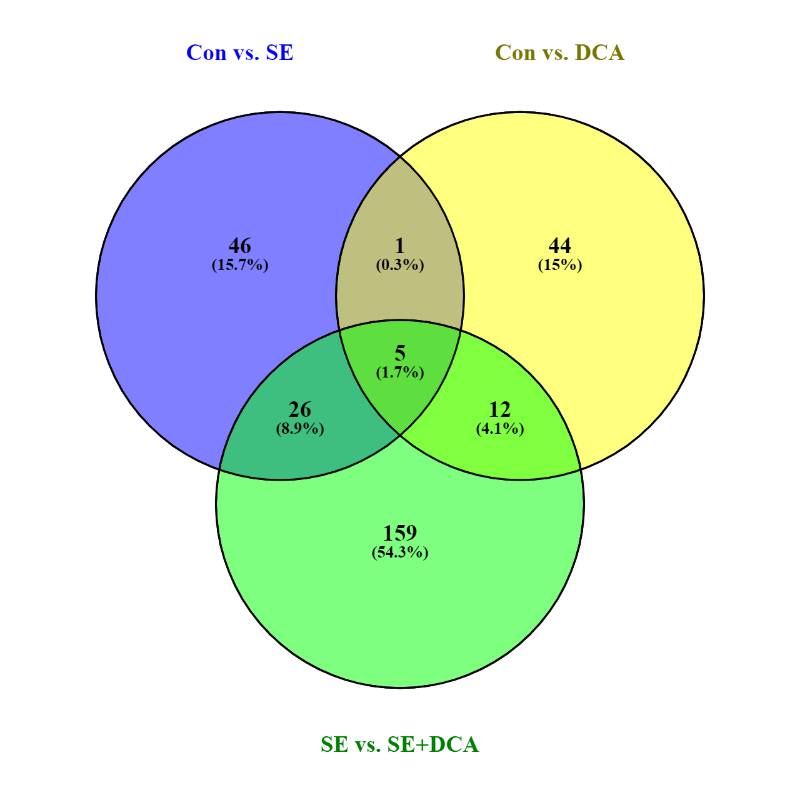

**Figure S3: Transcriptomic analysis of preterm pig cord blood following *S.epidermidis* stimulation and/or DCA supplementation. (A)** Principle component analysis plot based on gene expression profiles in treatment groups (n = 4/group). **(B)** Venn diagram demonstrating numbers of differentially expressed genes (DEGs) across group comparisons. Venn diagram was generated by venny 2.1 (<https://bioinfogp.cnb.csic.es/tools/venny/index.html>).

**Supplementary Tables:** uploaded separately due to the nature of big datasets
